# High intracellular calcium amounts inhibit activation-induced proliferation of mouse T cells

**DOI:** 10.1101/2024.04.02.587785

**Authors:** Joel P Joseph, Tanisha Kumar, Kaushik Chatterjee, Dipankar Nandi

## Abstract

Optimal T cell activation is critical to orchestrate adaptive immune responses. Calcium is critical for T cell activation and integrates signaling pathways necessary to activate key transcription factors. In fact, patients with calcium channelopathies are immunodeficient. Here, we investigated the effects of different concentrations of intracellular calcium on activation of mouse T cells. High intracellular calcium amounts inhibited *in vitro* T cell proliferation as evidenced by a decreased cell cycling-to-hypodiploidy ratio in two models of activation: the combination of phorbol 12-myristate 13-acetate (PMA) and Ionomycin (an ionophore)/Thapsigargin (a SERCA inhibitor) or plate bound anti-CD3 and anti-CD28. High intracellular calcium amounts increased the production of reactive oxygen species (ROS) in T cells activated with PMA and Ionomycin and scavenging excess ROS using N-acetyl cysteine (NAC) or PEGylated superoxide dismutase (PEG-SOD) rescued the decrease in cycling-to-hypodiploidy ratio. To test the universality of our observations, we studied the effects of tert-Butylhydroquinone (tBHQ), a SERCA inhibitor and Nrf2 activator. tBHQ alone did not increase intracellular calcium amounts but increase was observed along with PMA. Also, tBHQ inhibited T cell activation in a dose-dependent manner in both *in vitro* models of T cell activation. Importantly, intraperitoneal injection of tBHQ ameliorated Dextran Sodium Sulfate (DSS)-induced colitis in mice as evidenced by rescue of colon length shortening and lower disease activity index. Overall, this study identifies high calcium amounts as a potential target to lower T cell activation. The implications of these observations are discussed as this strategy may be important in treatment of some autoimmune diseases.

## INTRODUCTION

T cells play a critical role in adaptive immunity, where their optimal activation is key to eliciting an appropriate response. T cells recognize cognate antigens via the T cell receptors (TCRs), triggering a cascade of biochemical signalling pathways culminating in their activation, proliferation, and lineage differentiation (1). Calcium is a second messenger whose intracellular flux regulate transcription factors, such as nuclear factor of activated T cells (NFAT), nuclear factor kappa B (NF-κB), and activator protein 1 (AP-1), which activate the transcription of cytokine genes required for sustained T cell activation, proliferation, and differentiation (2–4). TCR stimulation generates inositol 1,4,5-trisphosphate (IP_3_), which opens the IP_3_ receptor channels and Ca^2+^ release from the endoplasmic reticulum. Calcium efflux from the endoplasmic reticulum activates STIM1, which then binds to ORAI1 and opens calcium release activated channels (CRAC), allowing calcium entry into the cytosol. Cytosolic calcium levels are further regulated by blocking calcium transport into the endoplasmic reticulum stores by inhibiting the sarcoendoplasmic reticulum calcium ATPase (SERCA) pump (5). On the one hand, the significance of intracellular calcium in T cell activation is clinically evidenced by immune deficiencies caused by calcium channelopathies, e.g., CRAC deficiency, which result in low intracellular calcium amounts (3). On the other hand, prolonged and aberrant T cell activation promotes chronic autoimmune diseases, such as multiple sclerosis, rheumatoid arthritis, systemic lupus erythematosus, etc (6). Previous reports have also shown that certain plant-derived compounds inhibit T cell activation by increasing intracellular calcium amounts. *Opuntia ficus indica* polyphenolic compounds (OFPC) increased calcium levels in a dose dependent manner but showed immunosuppressive effects on Jurkat T cells (7). A previous report from our laboratory showed that 7-hydroxyfrullanolide (7-HF) inhibits T cell activation by increasing intracellular calcium levels in mouse CD4^+^ T cells (8). However, a systematic investigation of the effects of higher amounts of intracellular calcium during T cell activation is lacking.

Furthermore, several drugs are used in the clinic to modulate calcium levels in different pathologies whose implication on the immune system is not well appreciated. For example, calcium channel blockers are routinely used in the treatment of various pathologies, such as hypertension, atherosclerosis, arrhythmia, etc (9). Calcium gluconate and other calcium supplements are administered to manage hypocalcemia (10). Thus, there is a need to understand how changes in intracellular calcium amounts play out in immunomodulation, specifically in T cell activation. Here, we investigated the direct effects of high amounts of intracellular calcium using known calcium ionophores with different modes of action, viz., Ionomycin and Thapsigargin. Ionomycin is a calcium ionophore (2), whereas Thapsigargin is a SERCA inhibitor (11). We evaluated T cell activation and associated proliferation with high intracellular calcium levels by estimating IL-2 release and cell cycling-to-hypodiploidy ratio of mouse primary T cells activated with different concentrations of the calcium ionophores. We also verified the principle using tert-butylhyroquinone (tBHQ), whose calcium-enhancing effects have been previously reported (11). Furthermore, we investigated the effects of tBHQ treatment in an autoimmune Dextran Sodium Sulfate (DSS)-induced colitis model in mouse.

## MATERIALS AND METHODS

### Animals

All experiments were performed using 6–8-week-old male C57BL/6 mice. For DSS treatment, mice were administered with 2.5% DSS (MP Biomedicals, USA) mixed in feeding water. At 48 and 96 h post DSS treatment, mice were intraperitoneally injected with tBHQ (Sigma-Aldrich, India). Mice were sacrificed on day 8 post treatment; colon length was measured, and blood was collected. All experiments were conducted as per the Control and Supervision rules 1998 of the Ministry of Environments and Forests Act, Government of India, and the Institutional Animal Ethics Committee of the Indian Institute of Science, under the permit number: CAF/Ethics/927/2022.

### T cell isolation

CD3^+^ T cells were isolated from inguinal, brachial, and mesenteric lymph nodes of 6–8-week-old, male, C57BL/6 mice, as previously described (12). Briefly, lymph nodes were mechanically disrupted and passed through a 70 µm cell strainer to prepare a single cell suspension. The cells were then panned in a T25 flask pre-coated with 100 μg/mL of AffiniPure goat anti-mouse IgG (Jackson Immuno Research laboratories, USA) to deplete the cell suspension of B cells, thereby enriching T cells. The purity of isolated T cells was estimated using flow cytometry (8,12).

### T cell activation

T cells were seeded at a density of ∼ 50,000 cells/well and activated using two models: (i) PMA (10 ng/mL) and Ionomcyin (0.1 or 0.5 μM) or Thapsigargin (0.01 or 0.05 μM), a TCR-independent activation model, to recapitulate optimal and high calcium levels (13,14) (ii) plate bound anti-mouse CD3 and soluble anti-mouse CD28, a TCR-dependent activation model (8,15). For anti-CD3/anti-CD28 based activation, 96 well U-bottomed plates (Corning, USA) were coated with 0.5 μg/mL of hamster anti-mouse CD3 (eBioscience, USA) and incubated overnight at 4°C (8). T cells were pretreated with PMA, Ionomycin, or tBHQ based on the treatment group and incubated at 37°C for 30 min before they were seeded on to anti-CD3-coated plates. Finally, 1 μg/mL of anti-mouse CD28 (eBioscience, USA) was added. Cell free culture supernatants were collected for ELISA, and cells were collected and fixed for cell cycle analysis at 36 h post activation.

### Cytokine measurements

IL2, IL6, TNFα, and IFNγ levels in cell-free culture supernatants or sera were estimated using eBioscience™ ELISA Ready-SET-Go™ kit as per manufacturer’s instructions (8). The standard curve was plotted by measuring the absorbance of appropriate dilutions (31.25 – 2000 pg/mL). The unknown cytokine amounts from the samples were interpolated from the respective standard curves within the detection range.

### Flow cytometry

T cells were stained for its surface markers by incubating the cells with pre-titrated dilutions (anti-mouse CD3 at 1:100; anti-mouse IgG at 1:200) of the conjugated antibodies. The cells were incubated in the dark for 45 min on ice with intermittent tapping. The cells are then washed with PBS and fixed with 0.5% paraformaldehyde in PBS (Sigma-Aldrich, USA). For cell size and cell cycle analyses, T cells were collected at ∼36 h post activation and fixed with 70% ethanol on ice for 30 min. The cells were then washed with 1× PBS and treated with 100 μg/mL of RNase A (Qiagen, USA) followed by staining with 50 μg/mL of propidium iodide (Sigma-Aldrich, USA). The cells were then incubated in the dark at room temperature for 30 min, and acquired on the flow cytometer (BD FACS Verse, BD Biosciences, USA). Data were analyzed using FloJo software. Single cell events were gated based on the forward scatter area versus forward scatter height, followed by gating with forward scatter area versus side scatter area. Histograms were generated and the median fluorescence intensity or percentage positive cells were estimated by marking the regions of interest.

### Intracellular calcium measurement

T cells were stained with 2 µM of Fluo-4 AM by incubating at 37°C in a humidified chamber for 1 h followed by activation using PMA and Ionomycin in the presence or absence of tBHQ. At 5 – 10 min post activation, cells were visualized using a confocal microscope (Andor Dragonfly, Nikon) with excitation/emission wavelength of 494/517 nm. Images were analyzed and intensity was measured using ImageJ software version 1.53t.

### Intracellular ROS measurement

T cells activated in the presence or absence of tBHQ were cultured for 36 h. At 36 h post activation, cells were collected, washed with 1× PBS and stained with 2′,7′-dichlorofluorescein diacetate (DCFDA, Sigma-Aldrich, USA) for 30 min at 37°C. Fluorescence intensity was measured using a microplate reader (TECAN, USA).

### DSS-induced colitis

Ulcerative colitis was induced in 6–8-week-old C57BL/6 mice using dextran sodium sulfate (DSS) by administering 2.5% (w/v) DSS in feeding water for seven days. Body weight, stool consistency, and hematochezia were recorded daily, and the disease activity index (DAI) was calculated as described previously (16). A body weight reduction of 20% was considered humane endpoint. To test the effect of tBHQ on DSS-induced colitis *in vivo*, mice were randomly divided into experimental groups (n = 3–7 each): control, colitis, and colitis + tBHQ treatment. The lead compounds were intraperitoneally injected at 5 mg/kg or 20 mg/kg on day 2 and day 4. On day 7, animals were sacrificed, and the endpoint analyses were performed. These included colon length measurement, H&E staining of tissues, and estimation of serum cytokine levels.

### Statistical Analysis

All statistical analyses and graphical representations were done on GraphPad Prism software version 8.0.2. Data are represented as mean ± standard error of mean (SEM) of three independent experiments, unless otherwise mentioned. Statistical analyses were performed using One-way ANOVA with Sidak’s test for multiple comparisons. *P*-value of less than 0.05 was considered significant, with statistical significance indicated as \**P* < 0.05, \*\**P* < 0.01, \*\*\**P* < 0.001, and \*\*\*\**P* < 0.0001.

## RESULTS

### High intracellular calcium inhibits T cell proliferation post activation with PMA and Ionomycin/Thapsigargin

To understand the effects of higher amounts of intracellular calcium on T cell activation and associated proliferation, primary mouse T cells were activated using PMA and different concentrations of ionomycin. First, we confirmed that ionomycin indeed increased intracellular calcium levels in a dose dependent manner (Figure 1a–b). The increase in calcium levels was more pronounced in the presence of PMA, suggesting the role of PKC activation in enhancing intracellular calcium levels in combination with a calcium ionophore, like Ionomycin, or enhanced calcium production by Ionomycin synergizing PKC activation (2). Next, we investigated morphological changes caused by PMA and ionomycin using a brightfield microscope. PMA or Ionomycin alone did not cause cell blasting; however, a combination of PMA and Ionomycin formed blast zones. There was an increase in cell blasting with increasing concentrations of ionomycin treatment in combination with PMA (Figure 1c), suggesting that high calcium increases T cell blasting. Furthermore, a combination of PMA and ionomycin increased IL-2 levels in a dose-dependent manner, suggesting increased IL-2 production with high intracellular calcium levels (Figure 1d). To investigate the effects of high amounts of intracellular calcium on activation-associated proliferation of T cells, we estimated the percentage of cycling and hypodiploid cells and their ratio using flow cytometry. The percentage of cycling cells increased and that of hypodiploid cells decreased with optimal calcium amounts i.e., at a combination of 10 ng/mL of PMA and 0.1 μM of Ionomycin. The cycling-to-hypodiploidy ratio also increased at this concentration. However, at a higher concentration of ionomycin, there was a decrease in the percentage of cycling cells, increase in the percentage of hypodiploid cells, and a consequent decrease in the cycling-to-hypodiploidy ratio (Figure 1e–h). These results suggested that high calcium levels decreased activation-associated proliferation in mouse T cells. To rule out the compound-specific effects of ionomycin and validate the principle independent of the mode of action of increasing cytosolic calcium levels, we performed similar experiments using a combination of PMA and Thapsigargin. Thapsigargin treatment also showed increased cell blasting and IL-2 release in a dose dependent manner, and decreased cycling-to-hypodiploidy ratio (Supplementary Figure 1). Furthermore, Ionomycin decreased the activation-associated increase in cell size; however, Thapsigargin treatment did not show any change in activation-associated cell size increase (Supplementary Figure 2).

**FIGURE 1.**
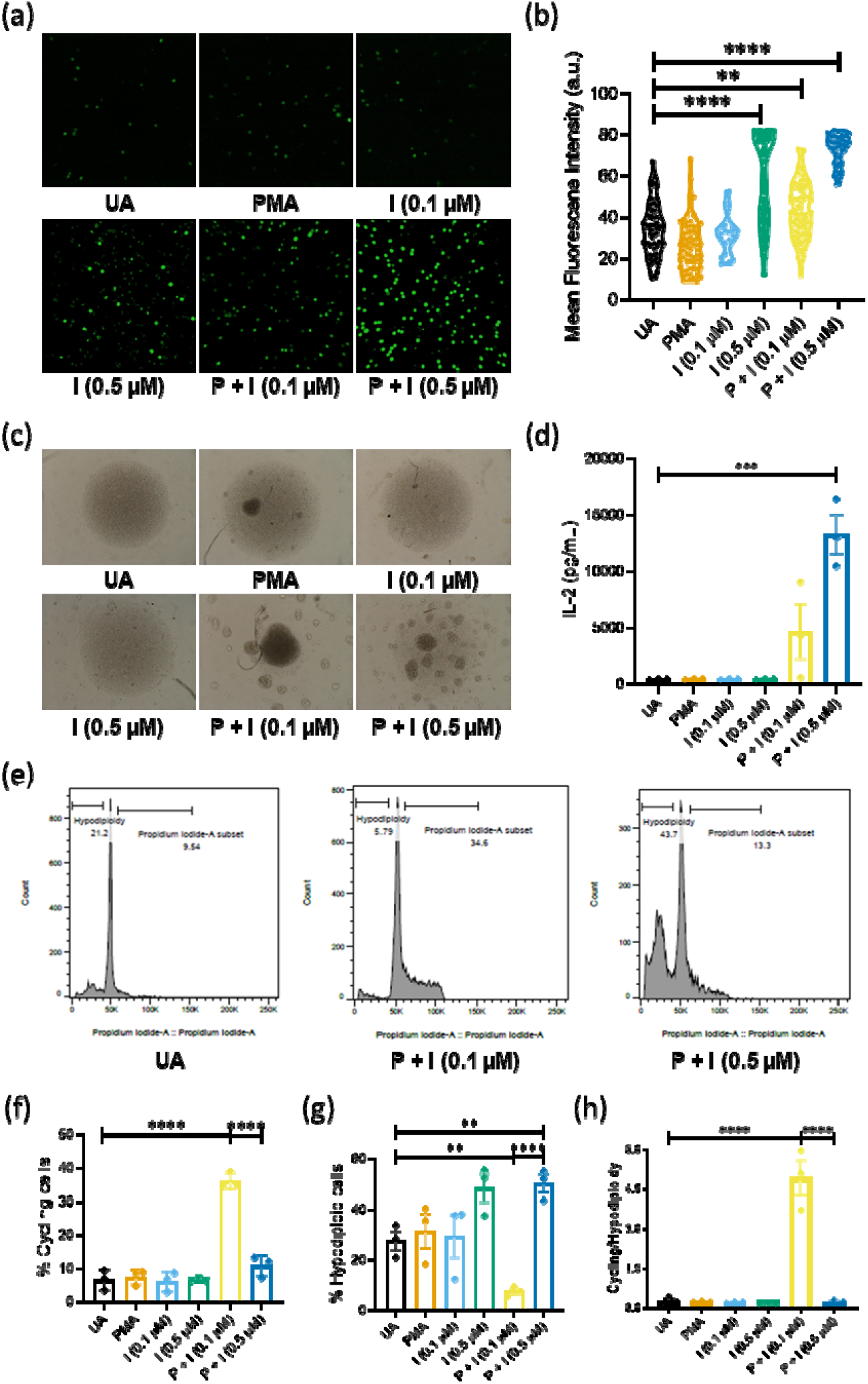
High intracellular calcium levels inhibit activation-associated proliferation in T cells in an *in vitro* T cell receptor independent activation model. (a) Fluo-4 AM staining of T cells treated with PMA and Ionomycin, (b) Mean fluorescence intensity of Fluo-4 AM staining, (c) Brightfield images of T cells treated with PMA and Ionomycin, (d) IL-2 amounts released into the supernatant of naive and activated T cell culture, (e) flow cytometric plots of cell cycling and hypodiploidy of T cell activated with optimal or higher amounts of intracellular calcium, (f) percentage of cycling cells in T cells activated with optimal or higher amounts of intracellular calcium, (g) percentage of hypodiploid cells in T cells activated with optimal or higher amounts of intracellular calcium, (h) cycling-to-hypodiploidy ratio. Data are represented as mean ± SEM from three independent experiments. For Fluo-4 AM staining, data are represented as mean ± SEM of individual cells from three independent experiments. One way ANOVA was performed to test the statistical significance, where \*\**p*<0.01, \*\*\**p*<0.005, \*\*\*\**p*<0.001. Abbreviations: P – PMA; I – Ionomycin.

### High calcium induces reactive oxygen species (ROS) production causing inhibition of activation-associated T cell proliferation

To decipher the mechanisms mediating high calcium induced inhibition of activation-associated T cell proliferation, we measured ROS production in cells using DCFDA staining. High intracellular calcium levels significantly increased ROS production in T cells (Figure 2a). Furthermore, scavenging ROS using N-acetyl cysteine (NAC) or Polyethylene Glycol conjugated Superoxide Dismutase (PEG-SOD) rescued the high calcium induced decrease in T cell cycling-to-hypodiploidy ratio (Figure 2b–c, Supplementary Figure 3), suggesting high calcium led to increased ROS production resulting in the inhibition of activation and associated increase in T cell proliferation.

**FIGURE 2.**
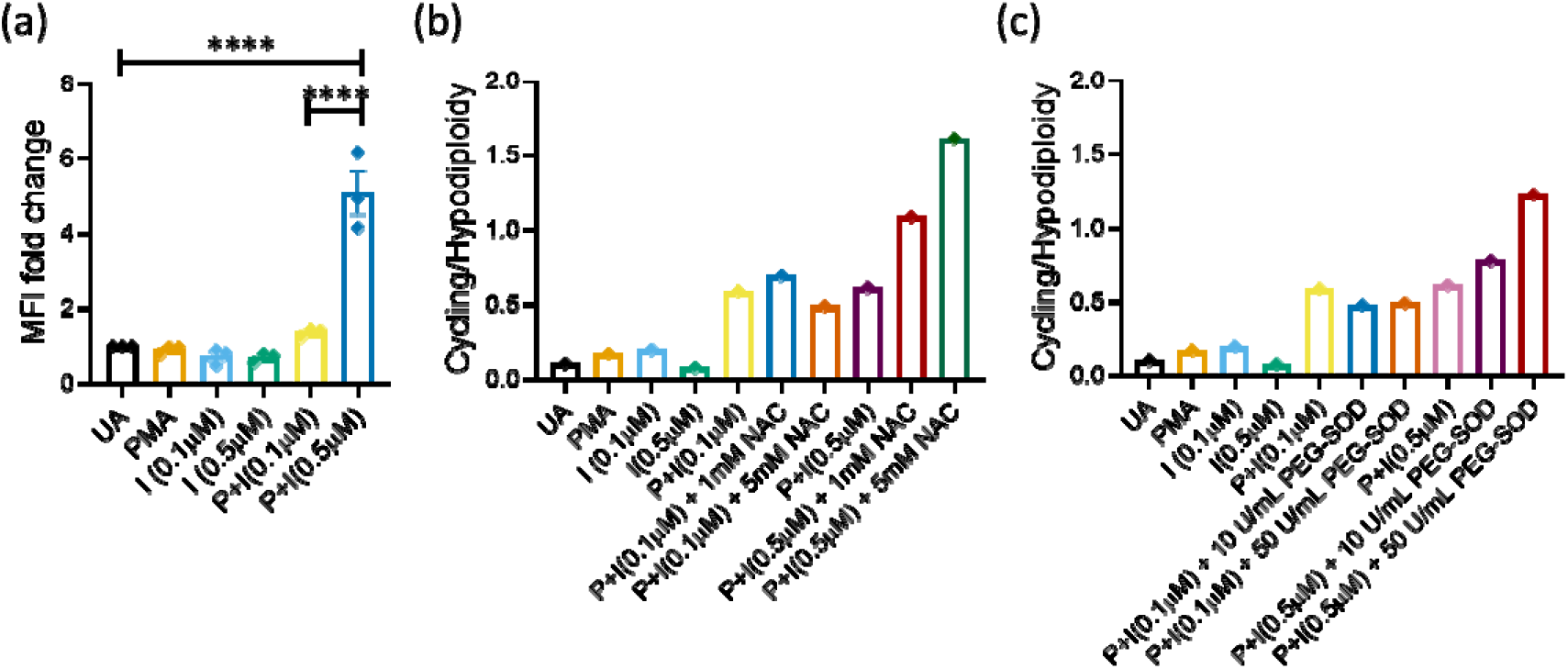
High intracellular calcium induces ROS production to inhibit activation-associated proliferation of T cells. (a) Relative fold change of mean fluorescence intensity upon DCFDA staining of T cells activated in optimal and high intracellular calcium levels, (b) cycling-to-hypodiploidy ratio of T cells activated in optimal and high intracellular calcium levels and treated with NAC, (c) cycling-to-hypodiploidy ratio of T cells activated in optimal and high intracellular calcium levels and treated with PEG-SOD. Data are represented as mean ± SEM from three independent experiments. One way ANOVA was performed to test the statistical significance, where \*\*\*\**p*<0.001. Abbreviations: NAC – N-acetyl cysteine; PEG-SOD – PEGylated super oxide dismutase; P – PMA; I – Ionomycin.

### High intracellular calcium levels inhibit T cell activation post activation with pb anti-CD3 and anti-CD28

To validate the findings in a T cell activation model that engages T cell receptor signaling, we activated T cells using plate bound anti-CD3 and anti-CD28 and treated them with PMA or Ionomycin. Activation of protein kinase C using increasing concentrations of PMA resulted in increased cell blasting upon anti-CD3 alone or anti-CD3 plus anti-CD28 condition (Figure 3a). Cycling-to-hypodiploidy ratio did not show a significant change although there was an increasing trend (Figure 3b). On the other hand, higher amounts of calcium inhibited activation-associated increase in cell blasting (Figure 3c) and decreased cycling-to-hypodiploidy ratio (Figure 3d), similar to our observation with the T cell activation model using PMA and Ionomycin. The IL-2 amounts in culture supernatants also significantly decreased with increasing activation of protein kinase C and increasing amounts of intracellular calcium (Supplementary Figure 5a–b).

**FIGURE 3.**
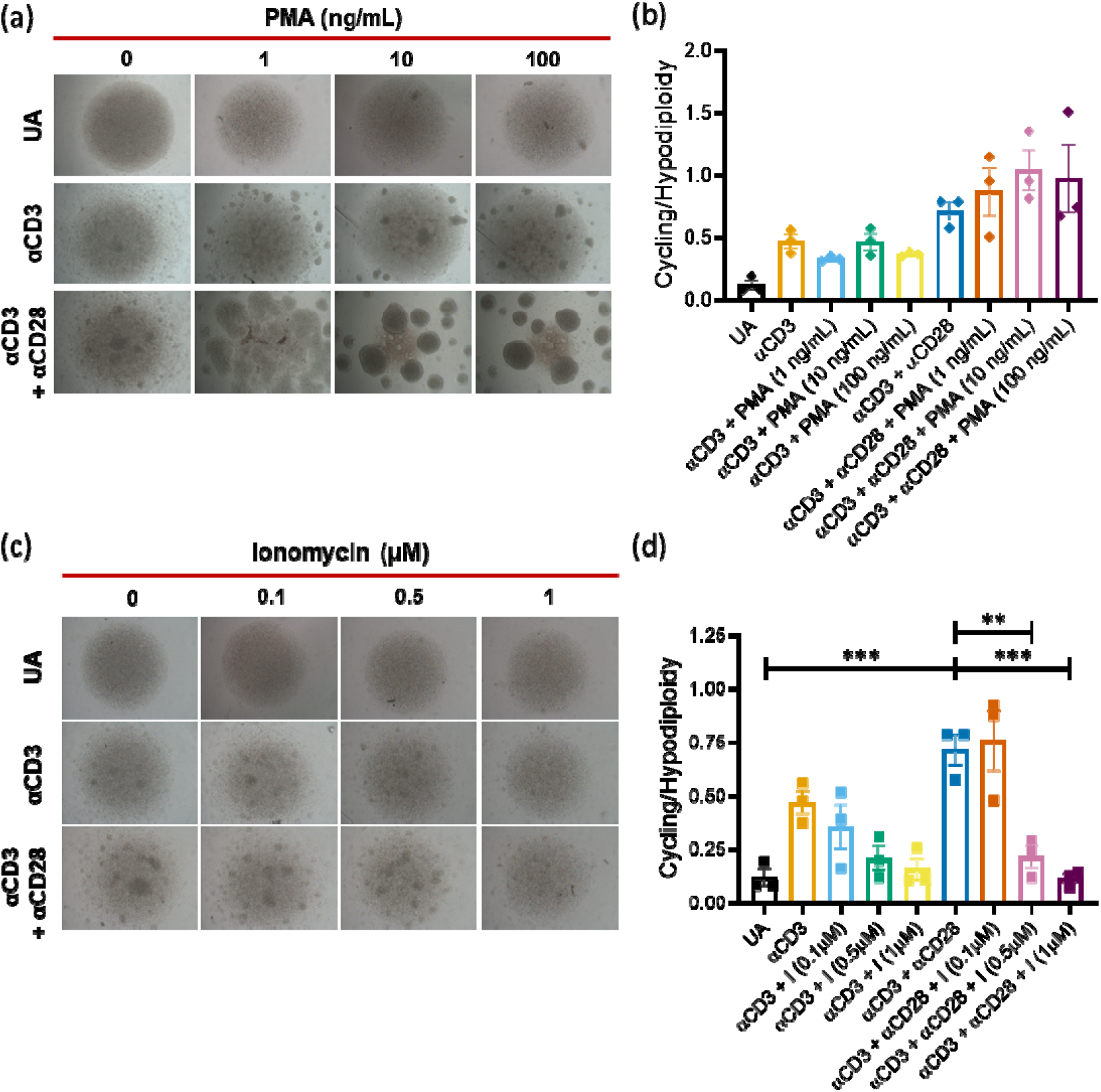
High intracellular calcium levels activation-associated proliferation in T cells in an *in vitro* TCR signaling dependent activation model. (a) Brightfield images of T cell activated with plate bound anti-CD3 and soluble anti-CD28 in the presence of increasing concentrations of PMA, (b) cycling-to-hypodiploidy ratio of T cell activated with plate bound anti-CD3 and soluble anti-CD28 in the presence of increasing concentrations of PMA, (c) brightfield images of T cells activated with plate bound anti-CD3 and soluble anti-CD28 in the presence of optimal and high intracellular calcium levels i.e., increasing concentrations of Ionomycin, (d) cycling-to-hypodiploidy ratio of T cells activated with plate bound anti-CD3 and soluble anti-CD28 in the presence of optimal and high intracellular calcium levels i.e., increasing concentrations of Ionomycin. Data are represented as mean ± SEM from three independent experiments. One way ANOVA was performed to test the statistical significance, where \*\**p*<0.01, \*\*\**p*<0.005.

### tBHQ increases intracellular calcium levels and inhibits T cell activation-associated proliferation

To understand whether the principle holds in the system using other calcium enhancers, we treated T cells with tBHQ, a Nrf2 activator (17,18) and SERCA inhibitor (11). In fact, the role of tBHQ as a calcium enhancer is reported in different cell types (11,19,20); however, its role in activation of primary mouse T cells is underappreciated. First, we tested whether tBHQ increases calcium levels in primary mouse T cells using Fluo 4-AM staining. While tBHQ alone did not increase calcium levels, it additively increased cytosolic calcium levels in combination with PMA and Ionomycin (Figure 4a–b). Furthermore, consistent with the effect of high amounts of calcium as displayed in increasing concentrations of Ionomycin with PMA, tBHQ decreased the cycling-to-hypodiploidy ratio in a dose dependent manner when treated above PMA and a low concentration of ionomycin (Figure 5a). tBHQ showed a similar effect T cell receptor dependent model of activation, where it decreased cell blasting and cycling-to-hypodiploidy ratio in a dose dependent manner (Figure 5b–c).

**FIGURE 4.**
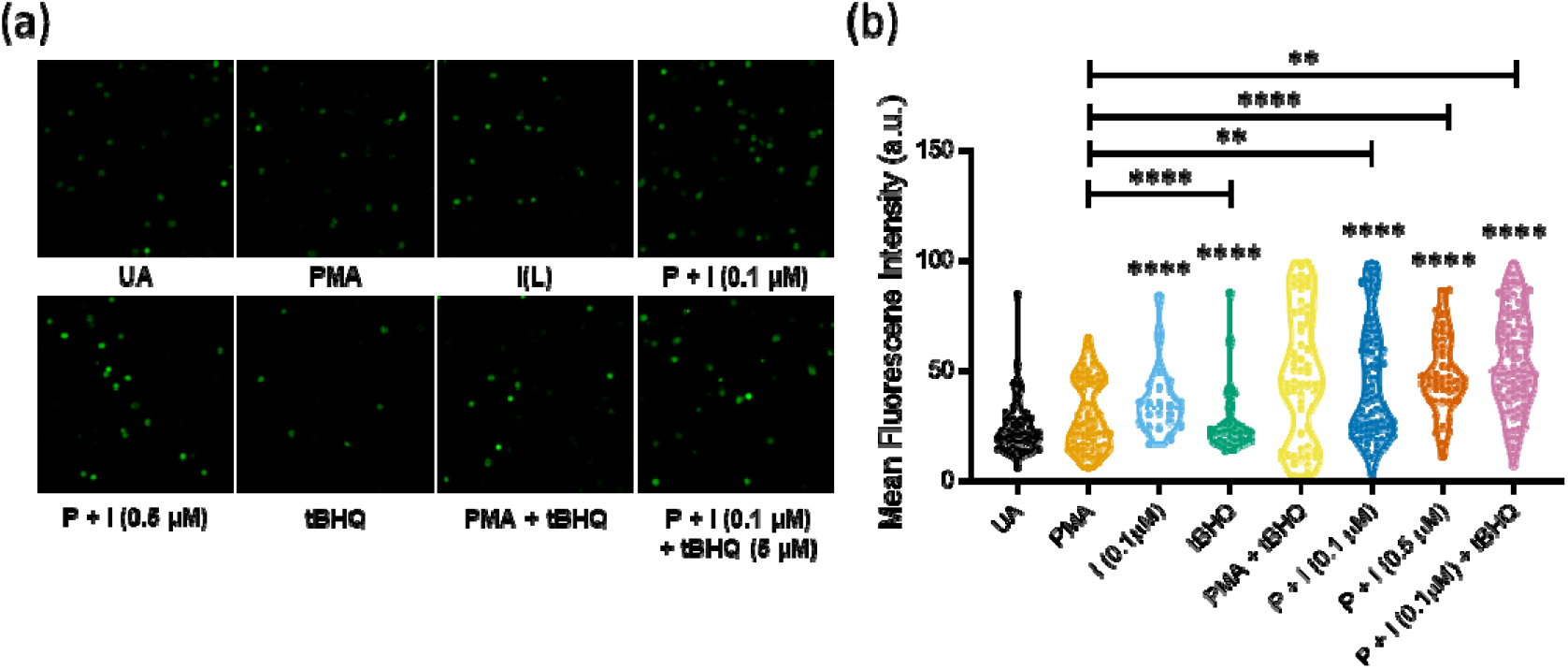
tBHQ increases intracellular calcium levels in an additive manner. (a) Fluo-4 AM staining of T cells treated with PMA and Ionomycin in the presence or absence of tBHQ, (b) mean fluorescence intensity of Fluo-4 AM staining. Data are represented as mean ± SEM of individual cells from three independent experiments. One way ANOVA was performed to test the statistical significance, where \*\**p*<0.01, \*\*\**p*<0.005, \*\*\*\**p*<0.001. Abbreviations: P – PMA; I – Ionomycin.

**FIGURE 5.**
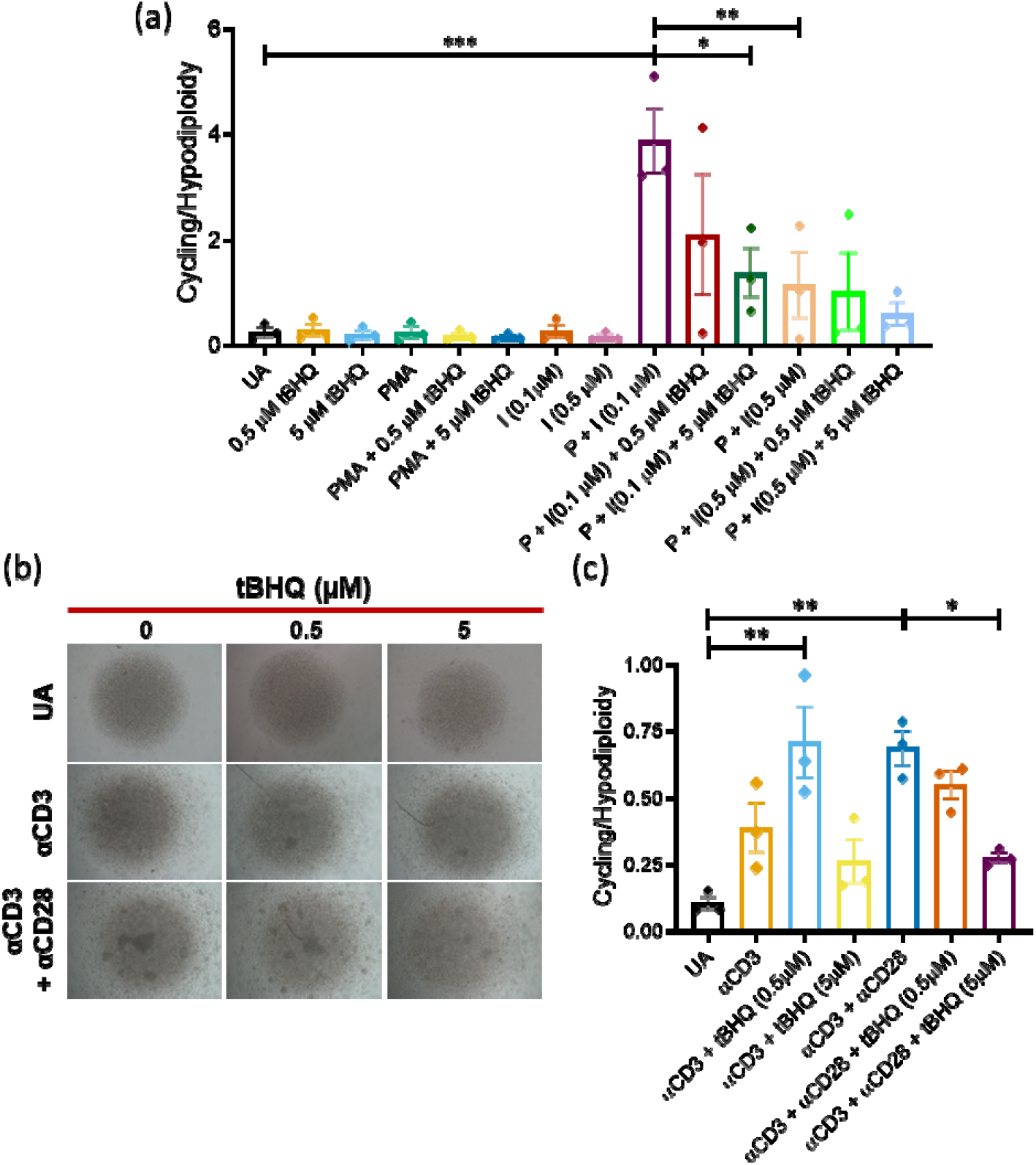
tBHQ inhibits T cell activation-associated proliferation in TCR independent and dependent activation models. (a) Cycling-to-hypodiploidy ratio of T cells activated with plate bound anti-CD3 and soluble anti-CD28 in the presence of optimal and high intracellular calcium levels i.e., increasing concentrations of tBHQ over PMA and Ionomycin. (b) brightfield images of T cells activated with plate bound anti-CD3 and soluble anti-CD28 in the presence of optimal and high intracellular calcium levels i.e., increasing concentrations of tBHQ, (c) cycling-to-hypodiploidy ratio of T cells activated with plate bound anti-CD3 and soluble anti-CD28 in the presence of optimal and high intracellular calcium levels i.e., increasing concentrations of tBHQ. Data are represented as mean ± SEM of individual cells from three independent experiments. One way ANOVA was performed to test the statistical significance, where \**p*<0.05, \*\**p*<0.01, \*\*\**p*<0.005.

### tBHQ ameliorates DSS-induced colitis in mice

To study the *in vivo* implications of high amounts of calcium in immune cell activation, we tested the effects of tBHQ in a mouse model of DSS-induced colitis. In DSS-induced colitis, immune cells, mainly T cells, macrophages and neutrophils, are aberrantly activated. We induced colitis in mice by administering 2.5% DSS in feeding water for 7 days (Figure 6a). In the treatment group, mice were intraperitoneally injected with two doses (5 mg/kg or 20 mg/kg) of tBHQ on day 2 and day 4 after commencing the DSS treatment. Mice injected with an equal dose of DMSO served as vehicle control. At low dose of 5 mg/kg body weight, tBHQ treatment did not produce any significant effect. However, at the higher dose of 20 mg/kg body weight, tBHQ significantly improved the colon length (Figure 6c–d), rescued disease activity index (Figure 6d), and decreased serum IL-6 levels (Figure 6e).

**FIGURE 6.**
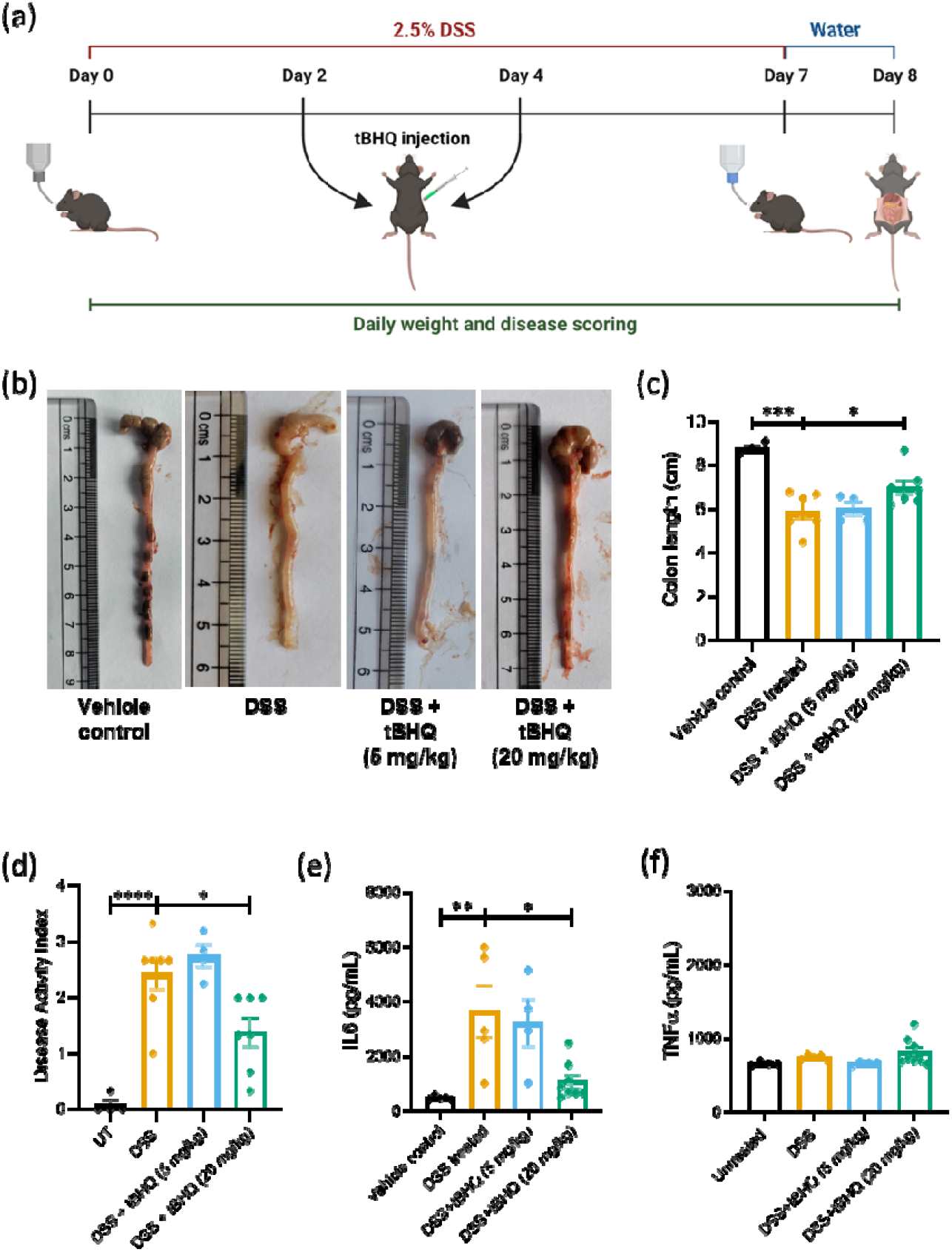
tBHQ rescued DSS induced colitis in mice. (a) Schematic of induction of DSS-induced colitis in mice and treatment with tBHQ, (b) Representative images of colons harvested from different experimental groups, (c) colon length of control, colitic and tBHQ treated mice, (d) disease activity index of control, colitic, and tBHQ treated groups, (e) serum IL6 levels in control, colitic, and tBHQ treated mice, (f) serum TNFα levels in control, colitic, and tBHQ treated mice. Data are represented as mean ± SEM from 3 – 7 mice per group. One way ANOVA was performed to test the statistical significance, where \**p*<0.05, \*\**p*<0.01, \*\*\**p*<0.005, \*\*\*\**p*<0.001.

## DISCUSSION

In this study, we investigated the direct effects of higher amounts of intracellular calcium on T cell activation using two compounds that increase cytosolic calcium levels: Ionomycin, a calcium ionophore (21), and Thapsigargin, a SERCA inhibitor (11,22). Also, we demonstrated that high intracellular calcium levels increased ROS production and hypodiploidy, and thereby inhibited activation-associated T cell proliferation. To further test the universality of the principle, we investigated the effects of tBHQ, an Nrf2 activator (18,23) and SERCA inhibitor (11) on *in vitro* mouse T cell activation. Although the role of tBHQ as SERCA inhibitor is appreciated in many cells, it is underappreciated in T cells. Moreover, we tested the effects of tBHQ treatment in an *in vivo* autoimmune disease model of DSS-induced colitis in mice (Figure 6).

Calcium influx is necessary for T cell activation, integrating pathways that result in the transcription of genes necessary for survival, proliferation, and differentiation. In fact, both TCR dependent and independent *in vitro* T cell activation models employ molecules that trigger calcium-modulated signaling pathway (3,13). Furthermore, T cells in mice and humans with calcium channelopathies have been reported to have impaired cytokine production (3,24). However, there is a lack of direct evidence for the effects of high intracellular calcium levels on T cell activation-associated events. A few reports have shown that plant-derived compounds could inhibit T cell activation by increasing intracellular calcium levels. For instance, 7-HF, a sesquiterpene lactone, inhibits T cell activation by increasing intracellular calcium levels in mouse CD4+ T cells (8). *Opuntia ficus indica* polyphenolic compounds increased calcium levels in a dose dependent manner but showed immunosuppressive effects on Jurkat T cells (7). However, it is unclear whether these effects are driven by other compound-specific cellular and molecular events or point to a universal principle correlating high intracellular calcium levels with inhibition of T cell activation. In this study, we found that high intracellular calcium levels increased cell blasting and IL-2 production in T cells activated with PMA and Ionomycin or Thapsigargin (Figure 1 & Supplementary Figure 1) but decreased them in a TCR dependent activation model that employed plate-bound anti-CD3 and anti-CD28 (Supplementary Figure 5b). In both the activation models, high calcium decreased the activation-associated proliferation of T cells by increasing ROS production in the TCR-independent model (Figure 2A, Supplementary Figure 3) and hypodiploidy in both activation models (Figure 1g, Supplementary Figure 4f).

Previous reports have shown that T cell activation induces ROS production, and that some amount of ROS is necessary for cell proliferation because ROS produced by respiratory complex I induces IL-2 and IL-4 expression in pre-activated human T cells (25). However, lower ROS production has also been associated with increased susceptibility to severe arthritis because of increased number of reduced thiol groups on T cell membrane surfaces (26). ROS production regulated surface redox levels of T cells and thereby suppressed autoreactivity and arthritis development (26), suggesting the importance of amounts of ROS production in regulating T cell activation.

In pathologies associated with other cells, such as neurons and muscles, it is evident that high calcium levels induce different types of cell death, including apoptosis by increased mitochondrial ROS production, and thus cause disease progression (27,28). Calcium dyshomeostasis and increased mitochondrial calcium levels in Alzheimer’s disease increase neuronal cell death and neurodegeneration in mice (29). High amounts of ROS and calcium also cause myofiber necrosis in muscular dystrophy (30,31). Our results demonstrated that high amounts of intracellular calcium produced higher amounts of ROS and decreased cell proliferation, as demonstrated in other cells, rather than further enhance T cell activation. As an *in vivo* proof-of-principle, we also showed that a calcium-enhancing drug with antioxidative properties, tBHQ, could ameliorate DSS-induced colitis in mice. A previous study investigating the effects of tBHQ on T cell activation in Jurkat cells reported a decrease in the percentage of cells positive for calcium influx, and a delayed calcium influx as assessed in the first 1.5 min after activation with anti-CD3/anti-CD28 (17). However, there was no inhibition of NFAT translocation or activation, concluding that the decrease in calcium influx may not have been great enough to diminish NFAT activation or that the decreased influx may have rebound at some later time point, ultimately allowing for full NFAT activation (17). In our observations with primary mouse T cells activated using PMA and ionomycin whose intracellular calcium levels were measured 5–10 min after activation, we found that tBHQ enhanced intracellular calcium levels when used in combination with PMA and Ionomycin. This small effect could be because the tBHQ sensitive calcium store present in the endoplasmic reticulum may be smaller than the one that has high affinity for other SERCA inhibitors, like Thapsigargin (11).

The study has several implications on the use of immunomodulatory drugs and calcium modulators. On one hand, calcium channel blockers are routinely used in the treatment of various pathologies, such as hypertension, atherosclerosis, arrhythmia, etc (9). Calcium signaling inhibitors are also used in treating immune pathologies, particularly autoimmune diseases. On the other hand, calcium gluconate and other calcium supplements are administered to manage hypocalcemia (10). Our observations imply that the use of such calcium enhancing drugs or direct administration of calcium could have implications on the immune system, such as inhibition of T cell activation-associated proliferation during an infection.

In conclusion, our study provides direct evidence that high intracellular calcium levels inhibit activation-associated proliferation of T cells by increasing ROS production and cell hypodiploidy. tBHQ, an Nrf2 activator and a calcium enhancer, inhibited activation-associated T cell proliferation *in vitro* and ameliorated DSS-induced colitis in mice. Our findings underscore the importance of dose-dependent responses and molecular details to immunomodulatory compounds that modulate intracellular calcium amounts. Further studies are needed to unravel the clinical implications of hypercalcemia and calcium enhancing drugs used in other treatments on T cell activation and T cell immune responses.

## AUTHOR CONTRIBUTIONS

JPJ performed experiments, analyzed data, and wrote the first draft of the manuscript; TK performed some experiments; KC supervised the study; DN conceptualized, designed, supervised the study, and revised the manuscript. All authors read and approved the manuscript.

## ACKNOWLEDGMENTS

This study was funded by core grants from IISc and the DBT-IISc partnership program. The authors gratefully acknowledge Dr Avik Chattopadhyay’s support with animal experiments and Dr Sanmoy Pathak’s mentorship to JPJ in training him with T cell activation models. We appreciate the support from members of the Central Animal Facility, IISc, Flow cytometry facility and Bio-imaging facility, Division of Biological Sciences, IISc. Thanks to Upasana Gupta for manuscript editing and review. The authors thank all members of DpN and KC laboratories for their support.

