## Supplemental Information for "High intracellular calcium amounts inhibit activation-induced proliferation of mouse T cells"

SUPPORTING INFORMATION

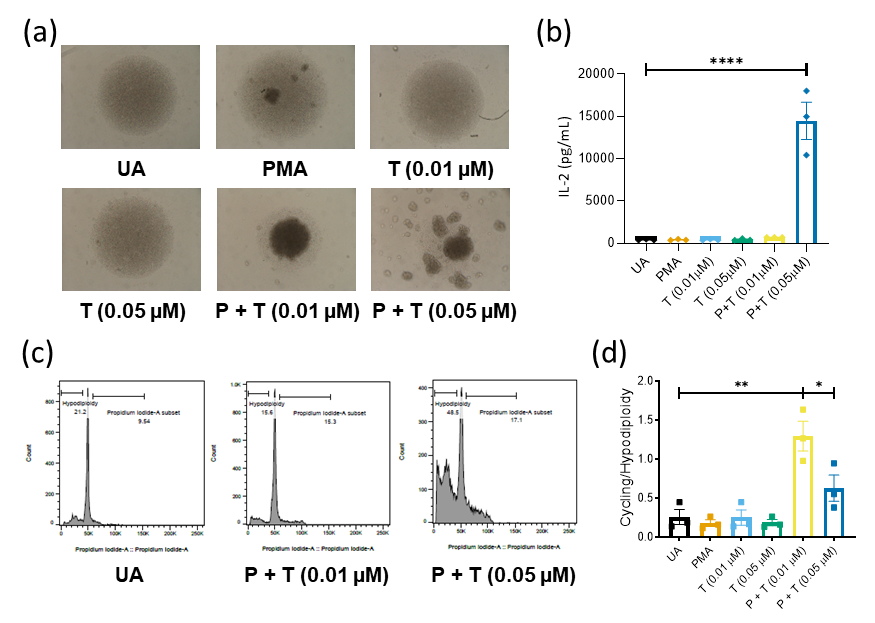

SUPPLEMENTARY FIGURE 1 High intracellular calcium levels inhibit activation associated proliferation in T cells in an *in vitro* T cell receptor independent activation model. (a) Brightfield images of T cells treated with PMA and Thapsigargin, (b) IL-2 amounts in culture supernatants of T cells treated with PMA and Thapsigargin, (c) flow cytometric plots of cell cycling and hypodiploidy of T cell activated with optimal or higher amounts of intracellular calcium i.e., different concentrations of Thapsigargin (d) cycling to hypodiploidy ratio of T cells activated with optimal or higher amounts of intracellular calcium. Data are represented as mean ± SEM from three independent experiments. One way ANOVA was performed to test the statistical significance, where **p*<0.05, ***p*<0.01, *****p*<0.001.

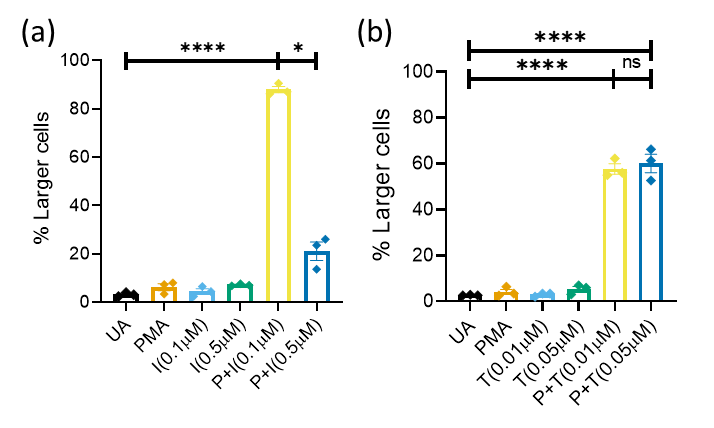

SUPPLEMENTARY FIGURE 2 High intracellular calcium levels induced by Ionomycin increases activation associated increase in T cell size. (a) Percentage of larger cells in T cells activated with PMA and different concentrations of Ionomycin recapitulating optimal and high intracellular calcium levels, (b) percentage of larger cells in T cells activated with PMA and different concentrations of Thapsigargin recapitulating optimal and high intracellular calcium levels. Data are represented as mean ± SEM from three independent experiments. One way ANOVA was performed to test the statistical significance, where **p*<0.05, *****p*<0.001.

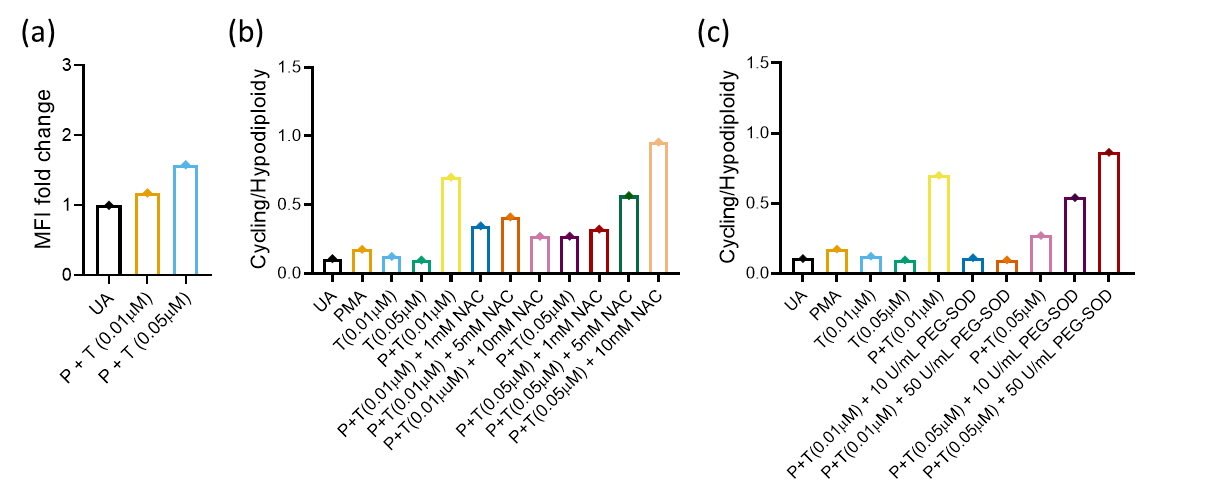

SUPPLEMENTARY FIGURE 3 High intracellular calcium induces ROS production to inhibit activation associated proliferation of T cells. (a) Relative fold change of mean fluorescence intensity upon DCFDA staining of T cells activated in optimal and high intracellular calcium levels, (b) cycling-to-hypodiploidy ratio of T cells activated in optimal and high intracellular calcium levels and treated with NAC, (c) cycling-to-hypodiploidy ratio of T cells activated in optimal and high intracellular calcium levels and treated with PEG-SOD.

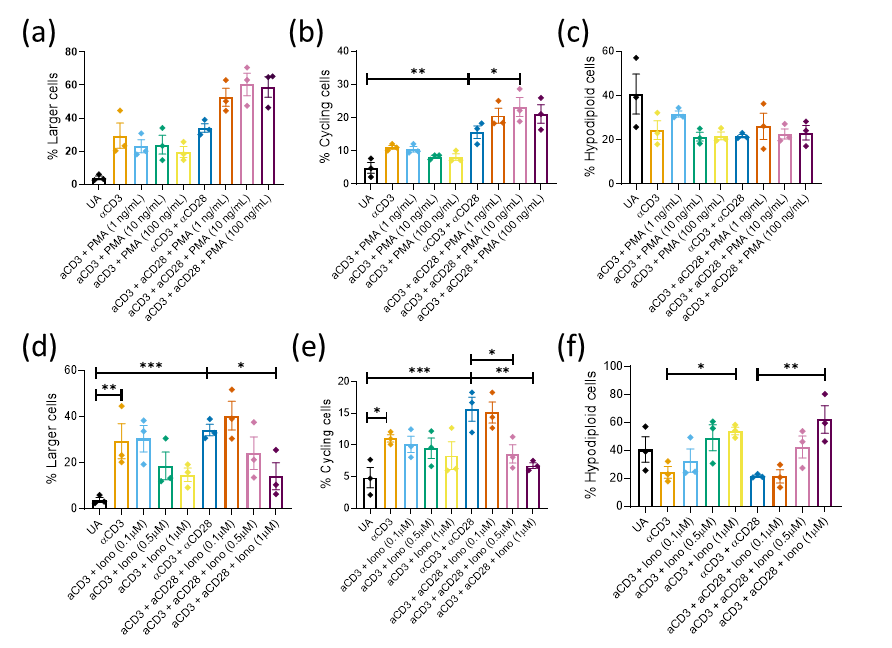

SUPPLEMENTARY FIGURE 4 Cell size, cycling, and hypodiploidy with high intracellular calcium levels in TCR-dependent activation model. (a) Percentage of larger cells with activation with anti-CD3 and anti-CD28 in the presence different concentrations of PMA, (b) percentage of cycling cells with activation with anti-CD3 and anti-CD28 in the presence different concentrations of PMA, (c) percentage of hypodiploid cells with activation with anti-CD3 and anti-CD28 in the presence different concentrations of PMA, (d) percentage of larger cells with activation with anti-CD3 and anti-CD28 in the presence different concentrations of Ionomycin, (e) percentage of cycling cells with activation with anti-CD3 and anti-CD28 in the presence different concentrations of Ionomycin, (c) percentage of hypodiploid cells with activation with anti-CD3 and anti-CD28 in the presence different concentrations of Ionomycin. Data are represented as mean ± SEM from three independent experiments. One way ANOVA was performed to test the statistical significance, where **p*<0.05, ***p*<0.01, *****p*<0.001.

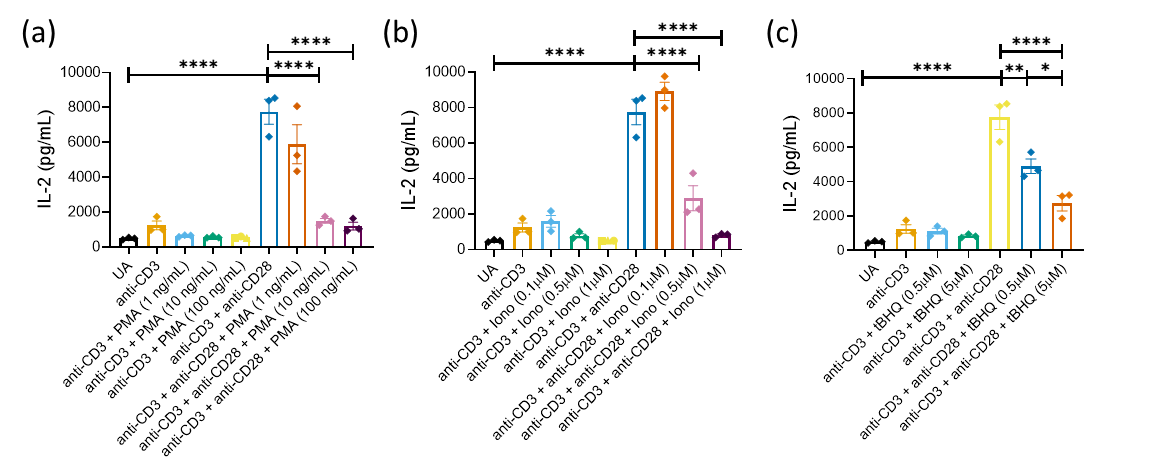

SUPPLEMENTARY FIGURE 5 IL-2 production upon high intracellular calcium levels in TCR dependent T cell activation. (a) IL-2 levels in culture supernatant of T cells activated with anti-CD3 and anti-CD28 in the presence of different concentrations of PMA, (b) IL-2 levels in culture supernatant of T cells activated with anti-CD3 and anti-CD28 and optimal and high intracellular calcium levels, i.e., different concentrations of Ionomycin, (c) IL-2 levels in culture supernatant of T cells activated with anti-CD3 and anti-CD28 in the presence of different concentrations of tBHQ. Data are represented as mean ± SEM from three independent experiments. One way ANOVA was performed to test the statistical significance, where **p*<0.05, ***p*<0.01, *****p*<0.001.
